# One invader, several origins: mitochondrial evidence of multiple introductions of *Rumina decollata* in Argentina

**DOI:** 10.64898/2026.02.22.707258

**Authors:** Julián Guerrero Spagnuoli, Néstor Sebastián Dop, Juan Manuel Rodrigo, Julia Pizá

## Abstract

Biological invasions comprise a major component of global change and biodiversity loss. Human-mediated dispersal introduced *Rumina decollata*, a Mediterranean land snail, beyond its native range, and the species is now widely distributed across much of Argentina. Here, we analysed mitochondrial COI sequences from specimens collected across a broad latitudinal and environmental gradient in Argentina to investigate their relationships with lineages reported from the native and introduced ranges. All analysed specimens clustered within the mitochondrial lineage previously identified as invasive worldwide. One haplotype identical to sequences reported from Spain and Portugal was detected in 21 of the 23 sampled localities, whereas two additional haplotypes matched lineages previously recorded from Portugal and southern France. These results support multiple introduction events into Argentina, followed by secondary spread. Body and sole colouration varied among individuals and showed no consistent association with mitochondrial lineages. Our findings provide the first broad-scale assessment of mitochondrial lineages of *R. decollata* in Argentina and contribute to understanding the introduction history of this invasive land snail.

## Introduction

Biological invasions comprise a major driver of global change and a leading cause of biodiversity loss (Bacher et al. 2024). Intentional or accidental introduction of invasive species beyond their native range results in the establishment of self-sustaining populations that can negatively impact native biodiversity, human health, economies, and cultural values (Machado and MacIsaac 2004; Essl et al. 2020). Understanding the pathways and origins of these invasions remains critical for developing effective management strategies, as climate change and global transport accelerate species introductions worldwide (Matheson and McGaughran 2022). In South America specifically, the number of documented non-native mollusc introductions and their impacts has increased markedly in recent years (Darrigran et al. 2025; Machado et al. 2026), highlighting the need for region-specific studies of invasion history within this group.

Land snails appear particularly prone to human-mediated dispersal via horticultural products, the pet trade, and cargo shipments (Cowie and Robinson 2003). While they play essential roles in ecosystems contributing to nutrient cycling, soil formation, and serving as prey for various organisms (Barker 2001), introduced land snails can outcompete or prey on native species, act as vectors for parasites, and cause economic losses in agriculture (Cowie et al. 2009). This makes understanding their introduction pathways and ecological impacts a priority in invasion biology.

Tracing the origins of invasive species often requires broad-scale genetic data (Puth and Post 2005; Facon et al. 2006). Analyzing haplotype variation in introduced populations and comparing it with reference sequences from native ranges can help to identify potential source regions and introduction routes (Puth and Post 2005; Facon et al. 2006; Prentis et al. 2008). Such approaches work particularly well for species with multiple introductions over large spatial and temporal scales.

*Rumina decollata* (Linnaeus, 1758) (Stylommatophora: Achatinidae), a terrestrial snail native to the Mediterranean region of southern Europe and northern Africa (Selander and Hudson 1976; Prévot et al. 2013a), has attained a widespread global distribution due largely to human-mediated, often accidental introductions (Prévot et al. 2014). As a hermaphroditic species capable of both cross- and self-fertilisation, it can reach high population densities, facilitating rapid colonization (Selander and Kaufman 1973; Prévot et al. 2014). Its xero-resistant nature allows it to inhabit dry, calcareous soils, waste grounds, disturbed habitats, and rocky terrains enabling survival in diverse environments and climates. Additionally, its omnivorous diet enables it to damage crops and outcompete or predate native fauna and flora (Tupen and Roth 2001; Matsukuma and Takeda 2009; Landal et al. 2019). *R. decollata* can also act as an intermediate host for parasites affecting rats and cats, with public health implications (Mas-Coma and Montoliu 1986; Cardillo et al. 2016, 2018).

Miquel (1988) first reported *R. decollata* in Buenos Aires province, Argentina, and it was subsequently recorded in several other provinces (De Francesco and Lagiglia 2007; Reyna and Gordillo 2018; Pérez and Tissot 2021; Rau et al. 2022). Recent surveys document its presence across urban and peri-urban areas spanning a wide range of climatic and environmental conditions (Pizá et al. 2023).

Early allozyme studies showed that the two previously described colour morphs of *Rumina decollata* correspond to distinct homozygous multilocus genotypes and differ in reproductive traits: the dark morph is associated with higher fecundity, whereas the light morph shows faster development, suggesting different life-history strategies with potential implications for colonization (Selander and Hudson 1976). Later, microsatellite analyses in native populations revealed high allelic diversity but low heterozygosity, consistent with frequent self-fertilisation (Lance et al. 2010), although higher heterozygosity in a native population from France suggests outcrossing may be more common than previously assumed (Prévot et al. 2013b). Experimental studies further indicate that, despite clear fitness costs associated with self-fertilisation, a considerable proportion of selfing individuals can successfully reproduce and produce viable offspring, highlighting the potential role of mixed mating systems in facilitating establishment after introduction (Pizá et al. 2025).

A molecular operational taxonomic unit (MOTU) is a cluster of individuals delimited by genetic similarity thresholds rather than by formal taxonomic description (Blaxter et al. 2005). MOTUs do not necessarily correspond one-to-one with morphologically defined species, and in facultatively self-fertilising taxa such as *Rumina* may represent cryptic or incipient lineages. Phylogenetic analyses of mitochondrial and nuclear DNA identified six such MOTUs within the native range of *Rumina decollata*. Among them, MOTU A (dark morph) includes all introduced populations examined to date in Asia and the Americas, whereas MOTU E comprises light-coloured individuals from the native range (Prévot et al. 2013a, 2014). Previous genetic studies of four Argentine localities suggested introductions from the Mediterranean region, particularly southern France and the Iberian Peninsula (Prévot et al. 2014; Rau et al. 2022). In addition, shell and sole colouration have historically been used to distinguish morphs within *R. decollata*, although the relationship between colour variation and molecular lineages remains incompletely understood.

To expand current knowledge of the introduction history of *R. decollata* in Argentina, we analysed mitochondrial COI sequences from specimens collected across a broad geographic and environmental gradient and compared them with previously published sequences from native and introduced populations. Specifically, we aimed to identify mitochondrial lineages present in Argentina, infer potential source regions, and evaluate whether colour morphs are consistently associated with the invasive lineage (MOTU A).

## Materials and methods

### Sample collection

Live snails of *Rumina decollata* were collected from 20 localities across 10 provinces of Argentina, covering diverse ecoregions, through a citizen science initiative (Pizá et al. 2023). Photographic verification of species identity confirmed a larger number of localities than the 20 included here (Pizá et al. 2023); the final set of sampling localities corresponds to the subset of photographically-verified records for which citizen scientists successfully collected and sent live specimens to our laboratory. No additional geographic stratification was applied. Snails were hand-collected by the citizen scientists at 19 of the 20 localities; specimens from Bahía Blanca (Buenos Aires Province) were collected directly by the authors. Snails were collected from private urban gardens, placed in plastic containers, and transported to our laboratory for further processing (Figure 1; Table S1).

**Figure 1.**
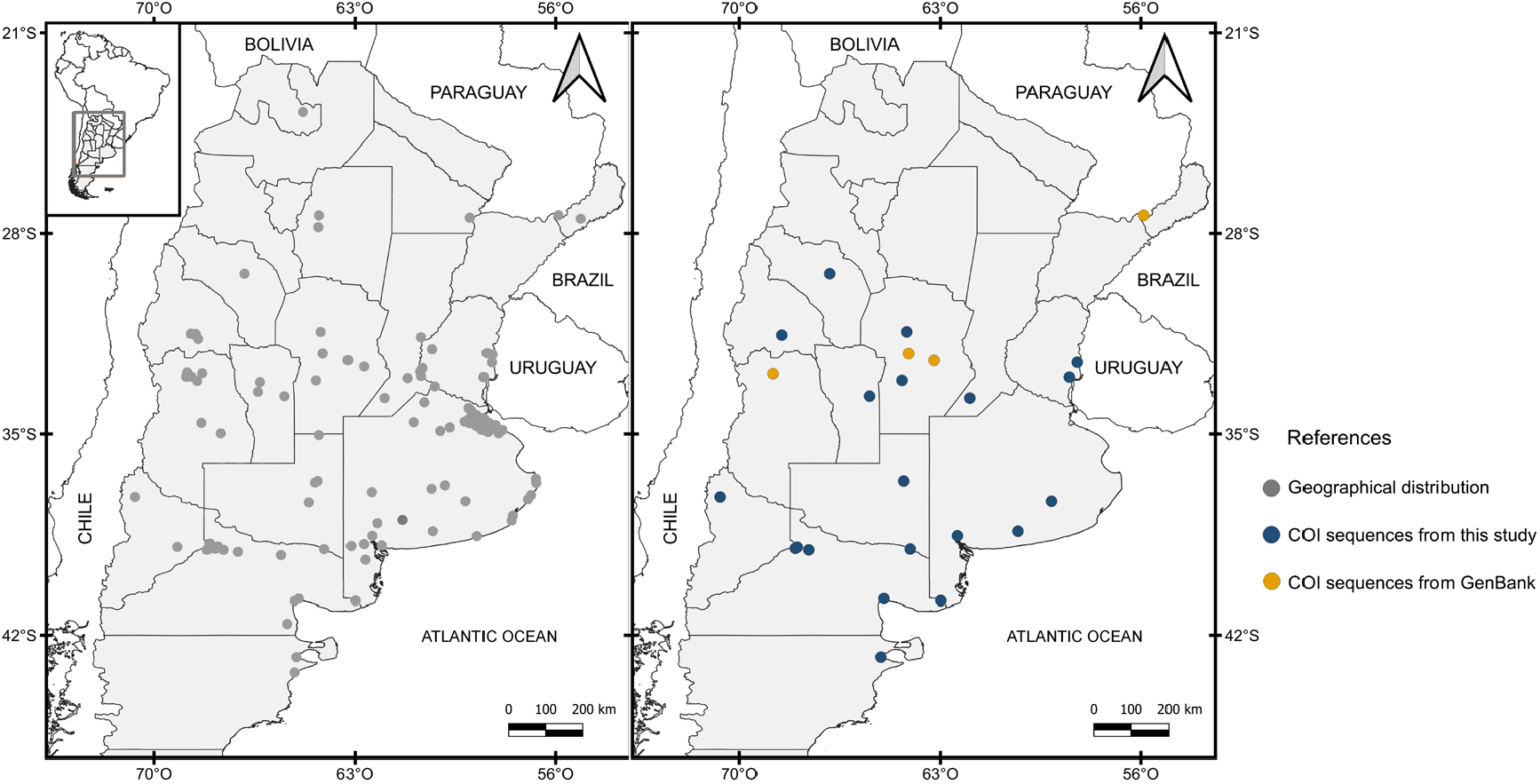
Geographical distribution and sample sources for *Rumina decollata*. Left: Map showing locations reported through the citizen science project (Pizá et al. 2023). Right: Distribution of samples used in genetic analysis. Blue dots: samples collected via the citizen science project. Orange dots: sequences available in GenBank.

## Genetic analysis

### Sample processing and DNA extraction

Snails were relaxed and sacrificed by immersion in deoxygenated water for approximately 12 h, following the technique used in previous anatomical studies of land snails by our group (e.g., Pizá and Cazzaniga 2009). Water was boiled to remove dissolved oxygen, sealed immediately, and allowed to cool to room temperature before snails were submerged; this method induces relaxation followed by death without triggering shell retraction. Snails were then preserved in 96% ethanol. Voucher specimens were deposited in the Malacological Collection of CERZOS (CONICET-UNS), Bahía Blanca, Buenos Aires Province, Argentina. Species identity was confirmed based on shell morphology (Prévot et al. 2015; Pizá et al. 2023).

DNA was extracted from a small piece of foot tissue from twenty individuals using an in-house CTAB-based protocol, adapted from standard CTAB extraction procedures commonly used for plant tissue (Doyle and Doyle 1987). We allowed ethanol to evaporate for 20 min, then incubated the tissue at 65°C for 30 min in 0.8 ml of CTAB buffer (2% [w/v] CTAB, 1.4 M NaCl, 0.2% [v/v] 2-mercaptoethanol, 20 mM EDTA, 100 mM Tris, pH 7.5). We purified DNA by triple chloroform extraction and precipitated it with cold 2/3 [v/v] isopropanol. The resulting DNA pellet was washed with 70% and 100% ethanol.

We successfully obtained DNA from 19 of the 20 samples collected; one sample did not yield amplifiable DNA and was excluded from further analyses. We selected one sample for each of the 19 sampled localities for genetic analysis.

### PCR amplification and sequencing

Partial sequences of the mitochondrial COI marker were amplified using primers LCO1490 and HCO2198 (Folmer et al. 1994). PCR master mix composition and thermal cycling followed Prévot et al. (2013a). Amplification products were verified via agarose gel electrophoresis. PCR products were bidirectionally sequenced by Macrogen Inc. (Seoul, Korea). Sequences were edited to remove primers and assembled into consensus sequences using BIOEDIT 7.2.5 (Hall 1999). Consensus sequences were compared to reference sequences from GenBank using BLASTn (Altschul et al. 1990) to support species identification. Sequences were aligned using ClustalW in Mega X (Kumar et al. 2018).

### Phylogenetic analysis

Phylogenetic relationships among haplotypes were inferred using the maximum likelihood (ML) method. All 23 Argentine sequences were included in the ML tree, rather than only the distinct haplotypes, in order to show the phylogenetic placement of every sampled locality individually and to allow direct comparison with the topology reported by Rau et al. (2022); the more compact set of distinct Argentine haplotypes is instead highlighted in the haplotype network (Figure 2). A total of 79 COI sequences were included: 23 from Argentina (19 new, 4 previously published; Prévot et al., 2014; Rau et al., 2022) and 56 from native and introduced populations retrieved from GenBank. *Subulina octona* (Bruguière, 1789) was used as an outgroup. ML analysis was conducted in Geneious Prime® 2022.1.1 using PhyML 3.0 (Guindon et al. 2010) with Nearest-Neighbor Interchange branch swapping, for consistency with the phylogenetic approach used in a previous analysis of Argentine *R. decollata* (Rau et al. 2022). The best-fit nucleotide substitution model (GTR+G) was selected with ModelFinder in IQTree (Kalyaanamoorthy et al., 2017) based on corrected AIC. Nodal support was assessed with 1,000 bootstrap replicates (Felsenstein 1985).

**Figure 2.**
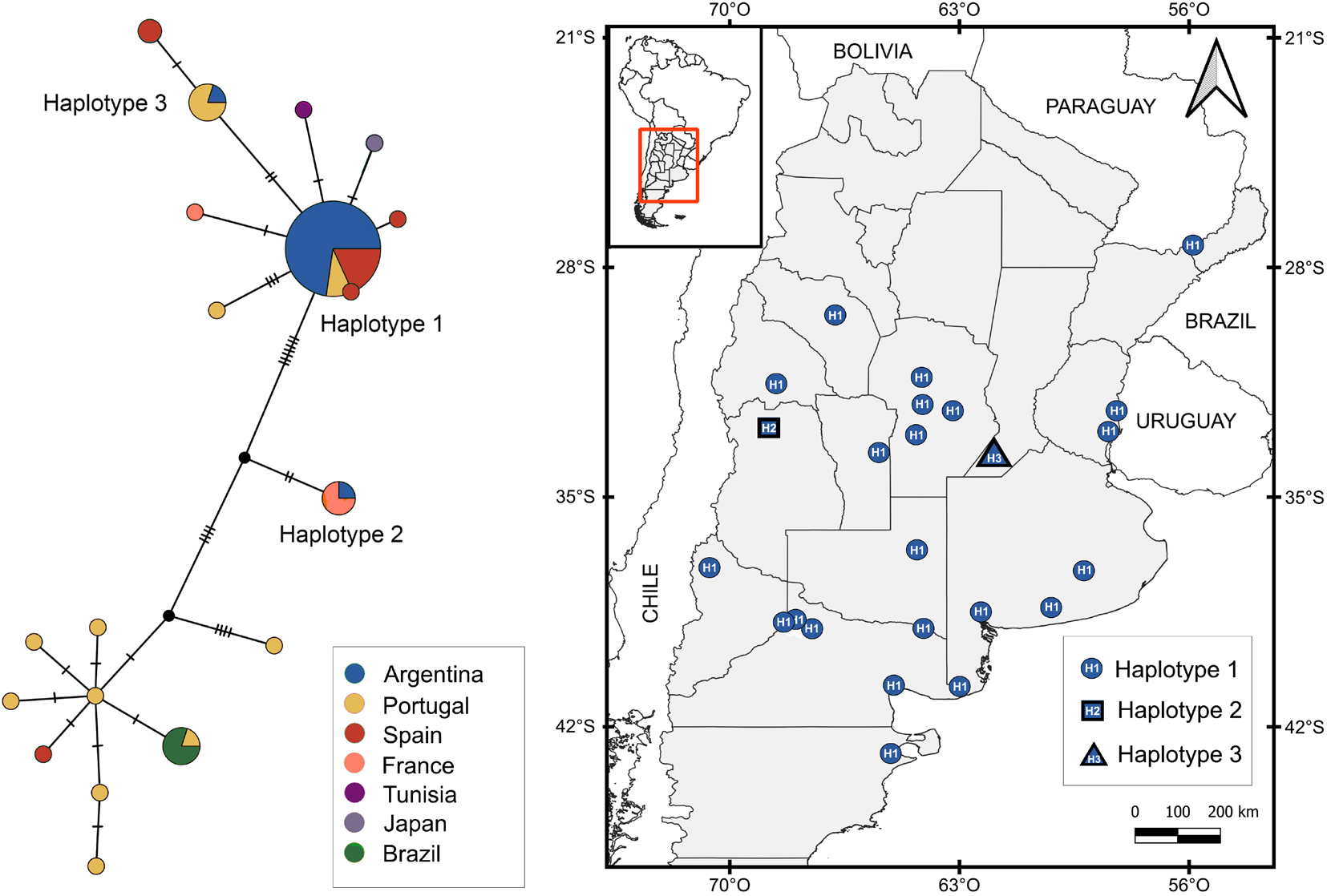
COI haplotype network and geographic distribution of *Rumina decollata* across the sampled localities in Argentina. Circle size in the network is proportional to haplotype frequency, and colours indicate the country of origin of reference sequences. Argentine haplotypes are labelled H1–H3, with H1 being the most frequent haplotype.

As all Argentine sequences grouped within MOTU A, a haplotype network was constructed using COI sequences from native and introduced populations assigned to this clade. The network was built with a median-joining algorithm (Bandelt et al. 1999) in PopART (Leigh and Bryant 2015). Sequences generated in this study were deposited in GenBank (accession numbers PP693485–PP693503).

### Morphological variability

To assess whether all individuals assigned to MOTU A correspond to the dark morph, we photographed live snails from each sampled locality and recorded shell and soft-part coloration (body and foot sole), following criteria described by Prévot et al. (2015).

## Results

### Source area of populations of *Rumina decollata* from Argentina

We obtained 19 COI gene sequences, each 655 base pairs in length. BLASTn searches supported the identification of all specimens as *R. decollata*. The sequences showed high similarity to those available in GenBank, with 100% coverage and top identity scores ranging from 99.85% to 100%.

Of the 23 Argentine sequences analysed (19 newly obtained and four retrieved from GenBank), only two showed genetic variation. One sample from Venado Tuerto (Santa Fe province) exhibited two nucleotide substitutions, while another sample from Mendoza province presented nine substitutions.

All *R. decollata* sequences from Argentina grouped within MOTU A in the phylogenetic analysis. The phylogenetic tree (Figure S1) and haplotype network (Figure 2) revealed three distinct haplotypes among the Argentine samples:six

1. Twenty-one of the 23 sequences matched haplotypes previously reported from populations in Spain and Portugal.
2. One sequence from Mendoza province matched a French haplotype (the “Arg” haplotype, Prévot et al. 2014).
3. One new haplotype from Venado Tuerto, Santa Fe province, matched a different haplotype from Portugal.

### Morphological variability

Body and foot sole coloration showed substantial inter-individual variation across the 20 sampled localities. Given this variability and the subjective nature of visual colour assessment, reliable assignment of individuals to previously described colour morphs was not possible (Figure 3).

**Figure 3.**
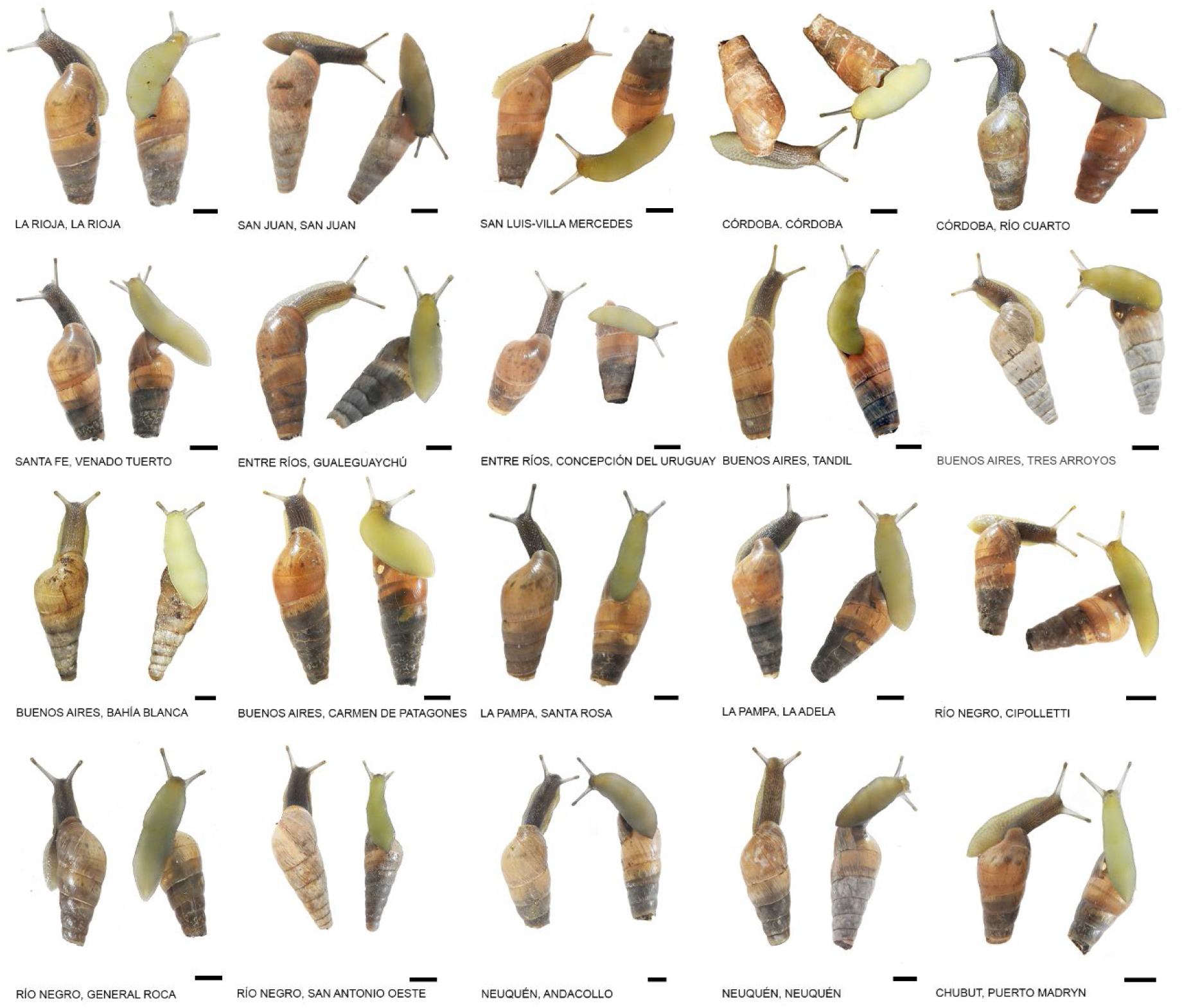
Representative individuals from the sampled localities of *Rumina decollata* illustrating variation in body and sole colouration. Scale bar: 5 mm.

## Discussion

This study provides the first broad-scale analysis of *Rumina decollata* in Argentina covering a wide biogeographical and latitudinal gradient, from Patagonia (42°45’S) to the Paranaense Forest (27°21’S). All Argentine populations clustered within mitochondrial molecular operational taxonomic unit A (MOTU A), previously identified as the lineage associated with introduced and colonizing populations worldwide (Prévot et al. 2014), reinforcing the hypothesis that only this lineage has successfully established outside the native range.

Our results are consistent with previous evidence for multiple introductions of *R. decollata* into Argentina, involving at least two source regions: the Iberian Peninsula and southern France (Prévot et al. 2014; Rau et al. 2022). A single Iberian haplotype, previously documented in Spain and Portugal (Prévot et al. 2013b), was detected in 21 of the 23 sampled localities, suggesting that this lineage represents the primary introduction source. In contrast, the sampled locality in Mendoza yielded a haplotype matching a sequence from southern France, whereas the locality of Venado Tuerto (Santa Fe) yielded a haplotype matching a sequence previously recorded from Portugal.

Taken together, these patterns suggest that the establishment of *R. decollata* in Argentina involved a limited number of independent introduction events, followed by extensive secondary spread within the country. Although the precise introduction pathways remain uncertain, accidental human-mediated transport via ornamental and horticultural plants is a plausible mechanism, as previously documented for land snails (Cowie and Robinson 2003). The species also occurs in Uruguay (Scarabino 2003) and southern Brazil (Landal et al. 2019). While no genetic data are available for Uruguay, Brazilian haplotypes cluster with Portuguese sequences that differ by 13 substitutions with Argentine ones, consistent with independent introductions into each country (Rau et al. 2022).

Despite the predominance of a single haplotype, the detection of three distinct haplotypes among the sampled localities (two of Iberian origin and one of French origin) indicates that introductions were multiple rather than the result of a single event. Repeated introductions from different source regions are often predicted to increase the genetic diversity of founding populations and may buffer the effects of demographic bottlenecks (Dlugosch and Parker 2008; Facon et al. 2008). However, our analysis, based on a single individual per locality, does not allow us to characterise haplotype diversity within populations, and the actual number of introduction events, as well as the extent of within-locality variation, may be underestimated. Analyses based on a larger number of individuals per site and on nuclear markers will be necessary to assess whether cryptic genetic structure or admixture has occurred.

The limited number of haplotypes recorded across the sampled localities is nevertheless consistent with expectations for invasive species that undergo strong founder effects during colonization (Dlugosch and Parker 2008; Charlesworth and Willis 2009), although this interpretation should be treated as preliminary given the sampling scale of this study. Such processes may facilitate the rapid spread of a few successful genotypes, particularly in species capable of establishing from very small propagule numbers.

Despite this apparent genetic homogeneity across the sampled localities, *R. decollata* has successfully colonised a wide range of environments in Argentina. Facultative self-fertilisation, documented in this species (Selander and Kaufman 1973; Prévot et al. 2013b; Pizá et al. 2025), provides reproductive assurance and allows populations to arise from very few individuals, even though it reduces effective population size and recombination (Lounnas et al. 2017). Although reproductive traits were not tested in this study, this life-history strategy offers a plausible, testable explanation for how *R. decollata* could establish and spread despite the limited number of mitochondrial haplotypes detected, and merits direct investigation in Argentine populations.

Finally, our data do not support a consistent association between MOTU A and dark body or sole colouration. Instead, individuals from Argentine populations exhibited a wide range of colour patterns. Visual assessment of colour is inherently subjective (Davison et al. 2019), and colour expression may be influenced by environmental factors such as temperature and light, which are known to affect pigmentation in other gastropods (Cowie and Jones 1985; Cowie 1990; Köhler et al. 2021). In the absence of standardized and quantitative assessment protocols, body and sole colouration cannot be considered reliable diagnostic traits for identifying MOTU A, at least in the Argentine populations examined here.

Overall, our results show that *Rumina decollata* has successfully established and expanded across a wide range of environments in Argentina, despite the limited number of mitochondrial haplotypes detected across the sampled localities. This pattern is consistent with the combined effects of repeated human-mediated introductions and life-history traits, such as facultative self-fertilisation, that could provide reproductive assurance, although this study did not directly test reproductive traits. Our results support the idea that invasion success does not necessarily require high genetic diversity at the scale examined here, and may instead arise from demographic processes and biological traits that buffer founder effects. The absence of a consistent association between body or sole colouration and the invasive lineage further emphasizes the limited value of morphological traits for diagnosing invasive clades in this species. Together, these results highlight *Rumina decollata* as a useful system for exploring how introduction history and reproductive strategies interact to facilitate biological invasions, and underscore the need for denser within-locality sampling and nuclear markers in future work.

## Acknowledgements

We are deeply grateful to the citizen scientists who contributed samples to this study. NSD, JMR and JP are members of CONICET (Consejo Nacional de Investigaciones Científicas y Técnicas, Argentina). We thank Dr. Echenique for providing laboratory facilities and consumables at the GENeTyC Laboratory, CERZOS (UNS-CONICET). We also thank Drs. Alda and Bonel for helpful comments that improved the manuscript.

## Statements and Declarations

### Ethics approval

Ethical review and approval were waived for this study because it involved land snails (invertebrates), which are not covered by specific ethical regulations in Argentina.

### Data availability

The COI sequences generated in this study are available in GenBank under accession numbers PP693485– PP693503. All other data generated or analysed during this study are included in this published article and its supplementary information file.

### Author Contributions

JP, JGS and NSD conceived and designed the study. JP supervised the project, secured the funding, and wrote the first draft of the manuscript. JGS, NSD and JP coordinated sample collection through the citizen science network. JMR provided technical training in molecular methods. JGS, NSD and JMR performed DNA extractions and PCR amplification. JGS and NSD conducted the phylogenetic and haplotype network analyses. All authors reviewed and approved the final version of the manuscript.

