## Supplemental material for "One invader, several origins: mitochondrial evidence of multiple introductions of *Rumina decollata* in Argentina"

[1] Genética y Ecología Evolutiva, Centro de Recursos Naturales Renovables de la Zona Semiárida, CONICET-Universidad Nacional del Sur, Bahía Blanca, Argentina

[2] Departamento de Biología, Bioquímica y Farmacia, Universidad Nacional del Sur, Bahía Blanca, Argentina

[3] Centro de Recursos Naturales Renovables de la Zona Semiárida, CONICET-Universidad Nacional del Sur, Bahía Blanca, Argentina

[4] Departamento de Agronomía, Universidad Nacional del Sur, Bahía Blanca, Argentina

†These authors contributed equally to this work and are co-first authors.

**Table S1.** Samples of living snails of *Rumina decollata* in Argentina obtained through citizen science project. Coordinates are estimates. Uncertainty was calculated according to the point-radius method (Wieczorek et al. 2004).

| Sample | Province, City | Latitude | Longitude | Uncertainty (km) | GenBank Accession |
|---|---|---|---|---|---|
| 1 | La Rioja, La Rioja | 29.4135° S | 66.8565° W | 7.38 | PP693493 |
| 2 | San Juan, San Juan | 31.5349° S | 68.5384° W | 2.87 | PP693498 |
| 3 | San Luis, Villa Mercedes | 33.6767° S | 65.4584° W | 4.45 | PP693485 |
| 4 | Córdoba, Córdoba | 31.4220° S | 64.1835° W | 8.49 | - |
| 5 | Córdoba, Río Cuarto | 33.1232° S | 64.3491° W | 6.22 | PP693490 |
| 6 | Santa Fe, Venado Tuerto | 33.7456° S | 61.9690° W | 4.43 | PP693489 |
| 7 | Entre Ríos, Concepción del Uruguay | 32.4845° S | 58.2321° W | 2.86 | PP693502 |
| 8 | Entre Ríos, Gualeguaychú | 33.0078° S | 58.5112° W | 3.36 | PP693496 |
| 9 | Buenos Aires, Tandil | 37.3288° S | 59.1367° W | 4.57 | PP693503 |
| 10 | Buenos Aires, Tres Arroyos | 38.3792° S | 60.2813° W | 2.60 | PP693501 |
| 11 | Buenos Aires, Bahía Blanca | 38.5382° S | 62.4049° W | - | PP693497 |
| 12 | Buenos Aires, Carmen de Patagones | 40.8014° S | 62.9897° W | 2.36 | PP693495 |
| 13 | La Pampa, Santa Rosa | 36.6209° S | 64.2912° W | 4.32 | PP693491 |
| 14 | La Pampa, La Adela | 39.0029° S | 64.0631° W | - | PP693487 |
| 15 | Río Negro, Cipolletti | 38.9388° S | 67.9954° W | - | PP693492 |
| 16 | Río Negro, General Roca | 39.0326° S | 67.5900° W | 3.60 | PP693486 |
| 17 | Río Negro, San Antonio Oeste | 40.7334° S | 64.9536° W | 3.02 | PP693488 |
| 18 | Neuquén, Andacollo | 37.1797° S | 70.6698° W | 0.64 | PP693499 |
| 19 | Neuquén, Neuquén | 38.9775° S | 68.0625° W | - | PP693500 |
| 20 | Chubut, Puerto Madryn | 42.7637° S | 65.0348° W | 3.90 | PP693494 |

**Figure S1.** Phylogenetic relationships of *Rumina decollata*. Maximum Likelihood tree based on partial COI gene sequences. Capital letters (A–F) indicate MOTUs previously defined by Prévot et al. 2013a. GenBank accession numbers and country of origin are indicated for each sequence. Bootstrap support values ≥70% are shown. Scale bar represents the number of substitutions per site.

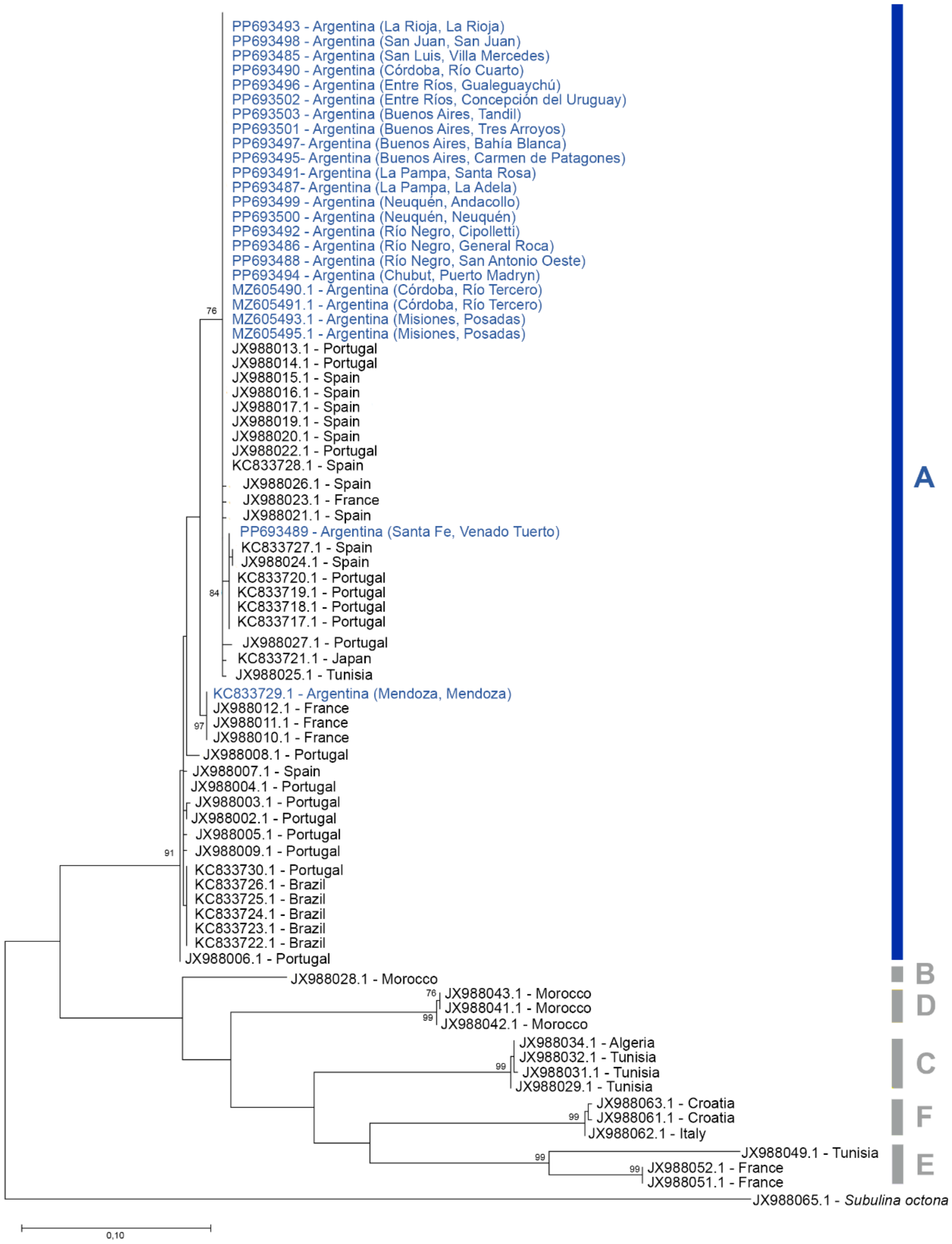
